## Supplementary material for "Phosphoproteomics identifies microglial Siglec-F inflammatory response during neurodegeneration": Key Resources Table

The table highlights the genetically modified organisms and strains, cell lines, reagents, software, and source data **essential** to reproduce results presented in the manuscript. Depending on the nature of the study, this may include standard laboratory materials (i.e., food chow for metabolism studies), but the Table is **not** meant to be comprehensive list of all materials and resources used (e.g., essential chemicals such as SDS, sucrose, or standard culture media don’t need to be listed in the Table). **Items in the Table must also be reported in the Method Details section within the context of their use.** The number of **primers and RNA sequences** that may be listed in the Table is restricted to no more than ten each. If there are more than ten primers or RNA sequences to report, please provide this information as a supplementary document and reference this file (e.g., See Table S1 for XX) in the Key Resources Table.

***Please note that ALL references cited in the Key Resources Table must be included in the References list.*** Please report the information as follows:

- **REAGENT or RESOURCE:** Provide full descriptive name of the item so that it can be identified and linked with its description in the manuscript (e.g., provide version number for software, host source for antibody, strain name). In the Experimental Models section, please include all models used in the paper and describe each line/strain as: model organism: name used for strain/line in paper: genotype. (i.e., Mouse: OXTR^fl/fl^: B6.129(SJL)-Oxtr^tm1.1Wsy/J^). In the Biological Samples section, please list all samples obtained from commercial sources or biological repositories. Please note that software mentioned in the Methods Details or Data and Software Availability section needs to be also included in the table. See the sample Table at the end of this document for examples of how to report reagents.
- **SOURCE:** Report the company, manufacturer, or individual that provided the item or where the item can obtained (e.g., stock center or repository). For materials distributed by Addgene, please cite the article describing the plasmid and include “Addgene” as part of the identifier. If an item is from another lab, please include the name of the principal investigator and a citation if it has been previously published. If the material is being reported for the first time in the current paper, please indicate as “this paper.” For software, please provide the company name if it is commercially available or cite the paper in which it has been initially described.
- **IDENTIFIER:** Include catalog numbers (entered in the column as “Cat#” followed by the number, e.g., Cat#3879S). Where available, please include unique entities such as [RRIDs](https://www.force11.org/group/resource-identification-initiative), Model Organism Database numbers, accession numbers, and PDB or CAS IDs. For antibodies, if applicable and available, please also include the lot number or clone identity. For software or data resources, please include the URL where the resource can be downloaded. Please ensure accuracy of the identifiers, as they are essential for generation of hyperlinks to external sources when available. Please see the Elsevier [list of Data Repositories](https://www.elsevier.com/authors/author-resources/research-data/data-base-linking) with automated bidirectional linking for details. When listing more than one identifier for the same item, use semicolons to separate them (e.g. Cat#3879S; RRID: AB_2255011). If an identifier is not available, please enter “N/A” in the column.
  - ***A NOTE ABOUT RRIDs:*** We highly recommend using RRIDs as the identifier (in particular for antibodies and organisms, but also for software tools and databases). For more details on how to obtain or generate an RRID for existing or newly generated resources, please [visit the RII](https://www.force11.org/group/resource-identification-initiative) or [search for RRIDs](https://scicrunch.org/resources).

Please use the empty table that follows to organize the information in the sections defined by the subheading, skipping sections not relevant to your study. Please do not add subheadings. To add a row, place the cursor at the end of the row above where you would like to add the row, just outside the right border of the table. Then press the ENTER key to add the row. Please delete empty rows. Each entry must be on a separate row; do not list multiple items in a single table cell. Please see the sample table at the end of this document for examples of how reagents should be cited.

***TABLE FOR AUTHOR TO COMPLETE***

*Please upload the completed table as a separate document.* ***Please do not add subheadings to the Key Resources Table.*** *If you wish to make an entry that does not fall into one of the subheadings below, please contact your handling editor. (****NOTE:*** *For authors publishing in Current Biology, please note that references within the KRT should be in numbered style, rather than Harvard.)*

**KEY RESOURCES TABLE**

| **REAGENT or RESOURCE** | **SOURCE** | **IDENTIFIER** |
| --- | --- | --- |
| **Antibodies** | | |
| Anti-phospho-Tyr (4G10) | Millipore | Cat# 05-321; RRID:AB_568857 |
| Anti-phospho-Tyr (PT66) | Sigma | Cat# P3300; RRID:AB_477335 |
| Anti-phospho-Ser/Thr-Pro (MPM-2) | Millipore | Cat# 05-368; RRID:AB_309698 |
| Anti-MAPK/CDK Substrate (34B2) | Cell Signaling Technology | Cat# 2325; RRID:AB_2797614 |
| Anti-Iba1 (Polyclonal) | Synaptic Systems | Cat# 234 004; RRID:AB_2493179 |
| Anti-Siglec-F (E50-2440) | BD Biosciences | Cat# 552125; RRID:AB_394340 |
| Anti-myc (9B11; Sepharose Bead Conjugate) | Cell Signaling Technology | Cat# 3400; RRID:AB_10692357 |
| Anti-myc (9B11) | Cell Signaling Technology | Cat# 9B11; RRID:AB_331783 |
| Anti-myc (71D10) | Cell Signaling Technology | Cat# 2278; RRID:AB_490778 |
| Anti-Siglec-8 (ab38578) | Abcam | Cat# ab38578; RRID:AB_777926 |
| Anti-Amyloid Beta (D54D2) | Cell Signaling Technology | Cat# 8243; RRID:AB_2797642 |
| Anti-MHC-II (HLA-DP, HLA-DQ, HLA-DR; CR3/43) | Agilent | Cat# M077501-2; RRID:AB_2313661 |
| Anti-SHP1 (C14H6) | Cell Signaling Technology | Cat# C14H6; RRID:AB_2173694 |
| Anti-SHP2 (D50F2) | Cell Signaling Technology | Cat# D50F2; RRID:AB_2174959 |
| APC-Cy7 Anti-Siglec-F (E50-2440) | BD Biosciences | Cat# 565527; RRID:AB_2732831 |
| APC Anti-CD33 (WM53) | BD Biosciences | Cat# 561817; RRID:AB_10896330 |
| Alexa-647 Anti-Siglec-5/14 (194128) | BD Biosciences | Cat# 564371; RRID: RRID:AB_2738773 |
| APC Anti-Siglec-8 (7C9) | Biolegend | Cat# 347105; RRID:AB_2561401 |
| Anti-IL-1β (Polyclonal) | R&D Systems | Cat# AF-401-SP;  RRID:AB_416684 |
| Anti-β-tubulin (Polyclonal) | Cell Signaling Technology | Cat# 2146, RRID:AB_2210545 |
| **Bacterial and Virus Strains** | | |
| Stbl3 | Thermo Fisher Scientific | Cat# C737303 |
| **Biological Samples** |  |  |
| Human AD brain samples | Banner Sun Health Research Institute | N/A |
| **Chemicals, Peptides, and Recombinant Proteins** | | |
| PBS | Thermo Fisher Scientific | Cat# 10010023 |
| Formaldehyde | Sigma | Cat# 252549 |
| Urea | Sigma | Cat# U5128 |
| Pierce BCA Protein Assay | Thermo Fisher Scientific | Cat# 23225 |
| Dithiothreitol (DTT) | Sigma | Cat# D0632 |
| Iodoacetamide (IAA) | Sigma | Cat# I1144 |
| Sequencing Grade Modified Trypsin | Promega | Cat# V5111 |
| 99.99% Acetic Acid | Sigma | Cat# 338826 |
| Sep-Pak Lite C18 Cartridge | Waters | Cat# WAT023501 |
| Sep-Pak Plus C18 Cartridge | Waters | Cat# WAT020515 |
| Pierce Quantitative Colorimetric Peptide Assay | Thermo Fisher Scientific | Cat# 23275 |
| 6-plex Tandem Mass Tag | Thermo Fisher Scientific | Cat# 90061 |
| 10-plex Tandem Mass Tag | Thermo Fisher Scientific | Cat# 90110 |
| ZORBAX 300Extend-C18 | Agilent | Cat# 770995-902 |
| High-Select™ Fe-NTA Phosphopeptide Enrichment Kit | Thermo Fisher Scientific | Cat# A32992 |
| 10 um C18 beads | YMC | Cat# ODS-A AA12S11 |
| 5 um C18 beads | YMC | Cat# ODS-AQ AQ12S05 |
| Poros 20 MC Metal Chelate Affinity Packing | Poros | Cat# 1-5429-06 |
| Fused Silica Capillary Tubing 50 μm ID | Polymicro Technologies | Cat# 1068150017 |
| Fused Silica Capillary Tubing 100 μm ID | Polymicro Technologies | Cat# 1068150023 |
| Fused Silica Capillary Tubing 200 μm ID | Polymicro Technologies | Cat# 1068150204 |
| Neuraminidase (Sialidase) from Arthrobacter ureafaciens | Sigma | Cat# 10269611001 |
| FuGene | Promega | Cat# E2311 |
| PEI MAX | Polysciences Inc | Cat# 24765-1 |
| .45 μm syringe filters | Sigma | Cat# 431220 |
| BD syringe | VWR | Cat# 302995 |
| RPMI 1640 Medium, GlutaMAX Supplement | Thermo Fisher Scientific | Cat# 61870036 |
| DMEM media | VWR | Cat# 10013CV |
| Fetal Bovine Serum (FBS), Certified | Thermo Fisher Scientific | Cat# 16000-044 |
| Pen Strep | Thermo Fisher Scientific | Cat# 15140-122 |
| 0.05% Trypsin | Thermo Fisher Scientific | Cat# 25300-120 |
| Annexin V binding/washing buffer | Thermo Fisher Scientific | Cat# V13246 |
| Annexin V-488 | Thermo Fisher Scientific | Cat# A13201 |
| Propidium Iodide | Thermo Fisher Scientific | Cat# P1304MP |
| PNGase F | New England BioLabs | Cat# P0704S |
| pHrodo Green Dextran | Thermo Fisher Scientific | Cat# P35368 |
| Superscript IV VILO Master Mix | Thermo Fisher Scientific | Cat# 11756050 |
| Ultrapure Water for HPLC | Sigma | Cat# 270733-4L |
| iQ SYBR Green Supermix | Biorad | Cat# 1708882 |
| Microseal 384-well PCR Plate | Biorad | Cat# MSP3842 |
| SHP099 | SelleckChem | Cat# S8278 |
| Bestatin | Sigma | Cat# B8385 |
| Tofacitinib citrate | Sigma | Cat# PZ0017 |
| Sodium Citrate | Sigma | Cat# S8750 |
| Hoechst 33342 | Thermo Fisher Scientific | Cat# H3570 |
| Matrigel | Corning | Cat# BD354277 |
| ReLeSR | StemCell Technologies | Cat# 05872 |
| mTESR | StemCell Technologies | Cat# 85850 |
| STEMdiff Hematopoietic Kit | StemCell Technologies | Cat# 05310 |
| ROCK inhibitor (Y-27632  dihydrochloride) | Tocris | Cat# 1254 |
| DMEM/F12 | Thermo Fisher Scientific | Cat# 11330-057 |
| ITS-G | Thermo Fisher Scientific | Cat# 41400045 |
| B27 | Thermo Fisher Scientific | Cat# 17-504-044 |
| N2 | Thermo Fisher Scientific | Cat# 17502048 |
| Glutamax | Thermo Fisher Scientific | Cat# 35050061 |
| MEM Non-essential Amino Acid (NEAA) Solution | Sigma | Cat# M7145-100ML |
| Penicillin:Streptomycin Solution | Gemini Bio-Products | Cat# 400-109 |
| Insulin | Sigma | Cat# 91077C-250MG |
| Monothioglycerol | Sigma | Cat# M6145-100ML |
| M-CSF | PeproTech | Cat# 100-21-1MG |
| IL-34 | PeproTech | Cat# 300-25-1MG |
| TGFβ-1 | PeproTech | Cat# 200-34-500UG |
| CD200 | Novoprotein | Cat# C311 |
| CX3CL1 | PeproTech | Cat# 300-31-1MG |
| Human IFN-α2 | StemCell Technologies | Cat# 78076 |
| Human IFN-β | R&D Systems | Cat# 8499-IF |
| Human IFN-γ | R&D Systems | Cat# 285-IF |
| Mouse IFN-γ | VWR | Cat# 575302-BL |
| TRIzol | Thermo Fisher Scientific | Cat# 15596018 |
| Chloroform | Sigma | Cat# C2432 |
| Ethanol, 200 Proof | VWR | Cat# V1001 |
| RIPA Buffer (2X) | Boston BioProducts | Cat# BP-115X |
| Halt Protease and Phosphatase Inhibitor Cocktail | Thermo Fisher Scientific | Cat# 1861281 |
| LDS Sample Buffer | Invitrogen | Cat# NP0007 |
| SDS-PAGE gel | Invitrogen | Cat# NP0335BOX |
| Nitrocellulose Membranes, 0.2 μm | Bio-Rad | Cat# 1620112 |
| Immun-Blot PVDF Membrane | Bio-Rad | Cat# 162-0177 |
| Novex Tris-Glycine Transfer Buffer | Thermo Fisher Scientific | Cat# LC3675 |
| Tris-Buffered Saline | Corning | Cat# 46-012-CM |
| Tween 20 | Fisher BioReagents | Cat# BP337 |
| Intercept Blocking Buffer | Li-Cor | Cat# 927-70001 |
| TrueBlack Lipofuscin Autofluorescence Quencher | Biotium | Cat# 23007 |
| **Critical Commercial Assays** | | |
| Direct-zol RNA MicroPrep Kit | Zymo Research | Cat# R2061 |
| Dynabeads mRNA Direct Kit | Thermo Fisher Scientific | Cat# 61012 |
| Kapa mRNA Hyperprep Kit | Roche | Cat# 08098115702 |
| pENTR/D-TOPO Cloning Kit | Thermo Fisher Scientific | Cat# K240020 |
| Gateway LR Clonase | Thermo Fisher Scientific | Cat# 11791020 |
| **Deposited Data** | | |
| BV-2 Siglec transcriptome | This paper | (https://www.ncbi.nlm.nih.gov/geo/) **GSEXXXX** |
| CK-p25, 5XFAD, and Tau P301S phosphoproteome | This paper | **PRIDE database (http://www.proteomexchange. org): PDX…** |
| **Experimental Models: Cell Lines** | | |
| HEK 293T cells | Forest White’s lab | ATCC CRL-3216; RRID:CVCL_0063 |
| BV-2 cells | Li-Huei Tsai’s lab | RRID:CVCL_0182 |
| Human induced-pluripotent stem cell line_AG09173 APOE3 | Li-Huei Tsai’s lab (Lin et al., 2018) | N/A |
| **Experimental Models: Organisms/Strains** | | |
| Mouse: CK-p25 | Li-Huei Tsai’s lab | Stock NO: 005706 |
| Mouse: 5xFAD: B6. SJL-Tg (APPSwFlLon, PSEN1*M146L*L286V)6799Vas/Mmjax) | Jackson laboratories | Stock NO: 34840 |
| Mouse: Δp35KI | Li-Huei Tsai’s lab | Stock NO: 022401 |
| Mouse: Tau P301S: B6;C3-Tg (Prnp-MAPT*P301S)PS19Vle/J | Jackson laboratories | Stock NO: 008169 |
| Mouse: C57BL/6 | Jackson laboratories | Stock NO: 664 |
| **Oligonucleotides** | | |
| Mouse IL-1B qPCR F:  GCAACTGTTCCTGAACTCAACT | Integrated DNA Technologies | N/A |
| Mouse IL-1B qPCR R:  ATCTTTTGGGGTCCGTCAACT | Integrated DNA Technologies | N/A |
| Mouse C1qa qPCR F:  AGGACTGAAGGGCGTGAAAG | Integrated DNA Technologies | N/A |
| Mouse C1qa qPCR R:  CGTGTGGTTCTGGTATGGACTC | Integrated DNA Technologies | N/A |
| Mouse GAPDH qPCR F:  TGGCAAAGTGGAGATTGTTGCC | Integrated DNA Technologies | N/A |
| Mouse GAPDH qPCR R:  AAGATGGTGATGGGCTTCCCG | Integrated DNA Technologies | N/A |
| Mouse B-actin qPCR F:  CTCTGGCTCCTAGCACCATGAAGA | Integrated DNA Technologies | N/A |
| Mouse B-actin qPCR R:  GTAAAACGCAGCTCAGTAACAGTCCG | Integrated DNA Technologies | N/A |
| Human Siglec-8 qPCR F: CAATATGGGGATGGTTACTTGCT | Integrated DNA Technologies | N/A |
| Human Siglec-8 qPCR R:  GGAGCGTCTTGGTATGGTCTG | Integrated DNA Technologies | N/A |
| Human GADPH qPCR F:  CCACCCATGGCAAATTCC | Integrated DNA Technologies | N/A |
| Human GADPH qPCR R:  TGGGATTTCCATTGATGACAA | Integrated DNA Technologies | N/A |
| Human B-actin qPCR F:  GGATGCAGAAGGAGATCACTG | Integrated DNA Technologies | N/A |
| Human B-actin qPCR R:  CGATCCACACGGAGTACTTG | Integrated DNA Technologies | N/A |
| **Recombinant DNA** | | |
| pCMV-VSV-G | Richard Hyne’s lab | N/A |
| pUMVC | Richard Hyne’s lab | N/A |
| psPAX2 | Addgene | Cat# 12260 |
| pMD2.G | Addgene | Cat# 12259 |
| MCSV-IRES-puro (MIP) | Richard Hynes’s lab | N/A |
| pINDUCER20 | Addgene | Cat# 44012 |
| MIP-Siglec-F-myc | This paper | N/A |
| MIP-Siglec-F-Y538F-myc | This paper | N/A |
| MIP-Siglec-F-Y561F-myc | This paper | N/A |
| MIP-Siglec-F-2xY->F-myc | This paper | N/A |
| MIP-hCD33-myc | This paper | N/A |
| MIP-hCD33-myc | This paper | N/A |
| MIP-hSiglec-5-myc | This paper | N/A |
| MIP-hSiglec-5-2xY->F-myc | This paper | N/A |
| MIP-hSiglec-8-myc | This paper | N/A |
| MIP-hSiglec-8-2xY->F-myc | This paper | N/A |
| pINDUCER20-Siglec-F-myc | This paper | N/A |
| pINDUCER20-Siglec-F-dVset-myc | This paper | N/A |
| pINDUCER20-Siglec-F-2xY->F-myc | This paper | N/A |
| pINDUCER20-hCD33-myc | This paper | N/A |
| pINDUCER20-hCD33-2xY->F-myc | This paper | N/A |
| pINDUCER20-hSiglec-5-myc | This paper | N/A |
| pINDUCER20-hSiglec-5-2xY->F-myc | This paper | N/A |
| pINDUCER20-hSiglec-8-myc | This paper | N/A |
| pINDUCER20-hSiglec-8-2xY->F-myc | This paper | N/A |
| **Software and Algorithms** | | |
| Proteome Discoverer (v2.2.1) | Thermo Fisher Scientific | <https://www.thermofisher.com/us/en/home/industrial/mass-spectrometry/liquid-chromatography-mass-spectrometry-lc-ms/lc-ms-software/multi-omics-data-analysis/proteome-discoverer-software.html> |
| MASCOT (v2.4.1) | Matrix Science | <http://www.matrixscience.com/> |
| XCalibur (v2.2) | Thermo Fisher Scientific | <https://www.thermofisher.com/order/catalog/product/OPTON-30965> |
| IncuCyte ZOOM (v2016A) | Sartorius | <https://www.essenbioscience.com/en/products/incucyte-zoom-resources-support/software-modules-incucyte-zoom/> |
| STAR (v2.5.3a) | (Dobin et al., 2013) | <https://github.com/alexdobin/STAR/releases> |
| RSEM (v1.3.0) | (Li & Dewey, 2011) | <http://deweylab.github.io/RSEM/> |
| DESeq2 (v1.18.1) | Bioconductor | <https://www.bioconductor.org/packages/devel/bioc/html/DESeq2.html>; RRID:SCR_016533 |
| GSEA (v3.0 beta-2) | (Mootha, et al., 2003; Tamayo et al., 2005) | <https://www.gsea-msigdb.org/gsea/> |
| R (v3.4.4) | N/A | <https://www.r-project.org/> |
| Salmon (v0.9.1) | (Patro, Duggal, Love, Irizarry, & Kingsford, 2017) | <https://combine-lab.github.io/salmon/> |
| Python (v3.7.1) | N/A | [https://www.python.org](https://www.python.org/) |
| MAGIC (v1.5.5) | (van Dijk et al., 2018) | <https://www.krishnaswamylab.org/projects/magic> |
| pyproteome (v0.11.0) | This publication | <https://github.com/white-lab/pyproteome> |
| scikit-learn (v0.21.1) | (Pedregosa et al., 2012) | <https://scikit-learn.org/stable/> |
| scipy (v1.3.1) | (Virtanen et al., 2020) | <https://www.scipy.org/> |
| numpy (v1.17.0) | (Van Der Walt, Colbert, & Varoquaux, 2011) | <https://numpy.org/> |
| pandas (v0.25.0) | (McKinney, 2010) | <https://pandas.pydata.org/> |
| seaborn (v0.10.0) | (Waskom, 2018) | <http://seaborn.pydata.org/> |
| matplotlib (v3.1.1) | (Hunter, 2007) | <http://matplotlib.org/> |
| goatools (v1.0.3) | (Klopfenstein et al., 2018) | <https://github.com/tanghaibao/goatools> |
| Cytoflow (v1.0) | N/A | <https://bpteague.github.io/cytoflow/> |
| MSigDB (v6.1) | (Liberzon et al., 2015; Subramanian et al., 2005) | <http://software.broadinstitute.org/gsea/msigdb/> |
| Fiji-ImageJ (v1.52u) | National Inst. Of Health | [https://imagej.net/Fiji](https://imagej.net/Fiji%20) RRID:SCR_003070 |
| ZEN (v2.1 SP3) | Zeiss | <https://www.zeiss.com/microscopy/us/products/microscope-software/zen.html> RRID:SCR_013672 |
| **Other** | | |
| Q Exactive Plus | Thermo | IQLAAEGAAPFALGMBDK |
| Q Exactive HF-X | Thermo | 0726042 |
| LTQ Orbitrap | Thermo | IQLAAEGAAPFADBMARX |
| VT100S vibratome | Leica | N/A |
| LSM 710 | Zeiss | N/A |
| LSM 880 | Zeiss | N/A |
| BD FACS Canto Cell Sorter | BD Biosciences | N/A |
| Li-Cor Odessey CLx | LI-COR Biosciences | N/A |
| Incucyte Plate Imager | Essen Bioscience | N/A |
| Illumina HiSeq 2000 | Illumina | N/A |
| CFX384 Touch Real-Time PCR | Biorad | N/A |
